# Bridging Higher-Order Information Theory and Neuroimaging: A Voxel-Wise O-Information Framework

**DOI:** 10.64898/2026.04.06.716652

**Authors:** Borja Camino-Pontes, Antonio Jimenez-Marin, Iñigo Tellaetxe-Elorriaga, Asier Erramuzpe, Ibai Diez, Paolo Bonifazi, Marilyn Gatica, Fernando E. Rosas, Daniele Marinazzo, Sebastiano Stramaglia, Jesus M. Cortes

## Abstract

The brain’s functional organization has been extensively studied through pairwise connectivity analyses. While these approaches have provided important insights into brain network organization, they fall short in capturing the complexity of high-order functional interactions (HOI). Particularly relevant is the investigation of redundancy and synergy patterns –not addressable with pairwise interactions–, revealing fundamental mechanisms of brain integration and information processing across various cognitive functions and clinical conditions. Conventional neuroimaging software packages are primarily designed for classical (general linear model-like) analyses but lack native support for HOI metrics. To address this gap, this study introduces a novel framework that bridges high-order information theory with conventional neuroimaging analysis pipelines and is subsequently applied to resting-state functional MRI to demonstrate its practical utility. By representing HOI into voxel-level metrics, our approach allows standard neuroimaging analyses to probe complex multivariate dependencies. Moreover, voxel-level group-comparison analyses show age differences linked with reduced redundancy in default mode network interactions. These findings advance our understanding of the complex relationship between multivariate functional interactions, voxel-level neuroimaging, and behavior, highlighting novel analytic strategies to study high-order information processing underlying cognitive function and its alterations in pathological conditions.

## Introduction

Understanding how different brain regions interact to support complex cognitive functions remains a fundamental challenge in neuroscience. Traditional connectivity analyses largely focus on pairwise relationships, measuring statistical dependencies such as correlations or mutual information between two brain areas at a time (Friston 1994; Bullmore and Sporns 2009; Meunier et al. 2010; Rubinov and Sporns 2011; Fornito et al. 2016). While these pairwise metrics have provided important insights into brain network organization, they fall short in capturing interactions among three or more brain regions simultaneously. Higher-order functional interactions play a crucial role in brain integration and information processing as shown in previous studies in healthy aging (Camino-Pontes et al. 2018; Gatica, Cofré, et al. 2021; Gatica, E. Rosas, et al. 2022), cognitive studies (Andrea I. Luppi et al. 2022), epilepsy (Erramuzpe et al. 2015; Mirjebreili et al. 2025), neurodegeneration (Herzog et al. 2022), physiology of population of neurons (Stramaglia et al. 2021), anesthesia (Andrea I Luppi et al. 2024), schizophrenia (Li et al. 2023), altered consciousness states (Herzog et al. 2022; Kumar G. et al. 2024) and transcranial ultrasound neuromodulation (Gatica, Atkinson-Clement, et al. 2025).

Despite the wide recognition of the importance of higher-order interactions, the majority of neuroimaging analysis tools—including widely used software packages such as SPM (Friston 2007), AFNI (Cox 1996), Nilearn (Abraham et al. 2014), CONN (Whitfield-Gabrieli and Nieto-Castanon 2012), fMRIprep (Esteban et al. 2019), and FSL (Jenkinson, Beckmann, et al. 2012) are primarily designed to perform classical general linear model (GLM) analyses in neuroimaging analysis. These tools do not natively support the integration of high-order interaction (HOI) metrics, limiting the possibility to investigate group differences or association with cognitive performance in patterns of redundancy and synergy at voxel- or region-level into GLM frameworks.

One of the main obstacles is the difficulty of visualizing and interpreting brain maps derived from interactions involving three or more variables. Unlike pairwise connectivity, where intuitive network diagrams and correlation matrices offer tangible representations, higher-order metrics require novel approaches to represent complex multivariate information into anatomically meaningful brain maps. This limitation has resulted in the emergence of two separated communities—conventional neuroimaging and HOI—highlighting the growing interest in tools capable of operating in both domains.

In this work, we introduce a novel yet powerful framework that, for the case of interactions within and between classical resting-state networks (RSNs) obtains maps of O-information (Rosas et al. 2019) susceptible to be analyzed by conventional neuroimaging. This approach aims to bridge the gap between the emerging field of higher-order information theory and the well-established neuroimaging analysis pipeline, facilitating the integration of multivariate interaction metrics into conventional workflows.

Specifically, we present four types of analyses that demonstrate the applicability and relevance of HOI mapping in neuroimaging. First, we generate population-level brain maps that highlight consistent patterns of redundancy and synergy across individuals. Second, voxel-level associations between these patterns and performance on standardized cognitive tests, revealing how higher-order dynamics relate to individual differences in cognition. Third, group differences in HOI maps between an old and a young group of participants. Finally, group differences between HOI and cognitive performance. Together, these analyses provide a comprehensive framework for integrating high-order metrics into the study of brain function and its variation across individuals and conditions.

## Methodology

### Neuroimaging Dataset

Neuroimaging data were obtained from the Max Planck Institute Leipzig Mind-Brain-Body Dataset (LEMON) (Babayan et al. 2018; Babayan et al. 2019), a rich multimodal dataset comprising MRI sequences, EEG, ECG, and behavioral scores. For this study, we focused on participants with both preprocessed fMRI and behavioral measures (N = 196). Our sample comprises healthy individuals aged 20–80 years (69 females; 35%). For detailed descriptions of these sequences, please refer to Table 7 in (Babayan et al. 2019).

### Behavioral Assessments

We selected a battery of neuropsychological tests targeting multiple cognitive domains relevant for diagnosing cognitive impairment in older participants: Regensburger Wortflüssigkeits-Test (RWT) (Aschenrenner et al. 2000), assessing verbal fluency and executive function; California Verbal Learning Test (CVLT) (Delis et al. 2016), assessing episodic memory and verbal learning; Trail Making Test parts A and B (TMT A and TMT B) (Reitan 1992), assessing attention and processing speed; and, from the Test of Attentional Performance (TAP) (Leclercq and Zimmermann 2002), alertness (TAP A), visual attention (TAP VIS), motor inhibition (TAP HAND), and working memory (TAP WM). For a detailed description of the psychological assessment, please refer to the Methods section in (Babayan et al. 2019). In general, higher scores indicate better performance. For TMT A, TMT B, TAP A, TAP VIS, and TAP HAND, lower scores indicate better performance, as they reflect shorter completion times or faster reaction times. These scores were inverted (multiplied by –1) before analysis to ensure that higher values consistently indicate better cognitive performance across all tests. From here on, inverted scores are denoted with an overline (i.e., TMT), indicating that higher values represent better performance.

### Brain Imaging Preprocessing

The functional HOIs were assessed from time-series data provided by LEMON. Although the pre-processing details can be found in (Mendes et al. 2019), major steps included: 3D motion correction (FSL MCFLIRT) (Jenkinson, Bannister, et al. 2002), distortion correction (FSL FUGUE) (Jenkinson, Beckmann, et al. 2012), rigid-body co-registration of the unwrapped temporal mean image to the individual’s anatomical image (FreeSurfer bbregister) (Greve and Fischl 2009), denoising (Nipype rapidart and aCompCor) (Behzadi et al. 2007), band-pass filtering between 0.01–0.1 Hz (FSL), mean centering, variance normalization of the denoised time series (Nitime) (Rokem et al. 2009), and spatial normalization to MNI152 at 2 mm resolution via transformation parameters derived during structural preprocessing (ANTs SyN) (Avants et al. 2011).

We further used two complementary tools to mitigate physiological confounds in the BOLD signal and improve the estimation of HOI. RapidTide (Frederick 2016-2024) was applied to detect lagged signal propagation patterns within the BOLD signal, reducing spurious correlations caused by systemic low-frequency oscillations (sLFOs) (Korponay et al. 2024). In addition, the resting-state hemodynamic response function (rsHRF) was estimated using a blind procedure based on point processes and a mixture of Gamma functions (Wu et al. 2021), and subsequently deconvolved to remove temporal confounds (Rangaprakash et al. 2018). These preprocessing steps ensure that the voxel-level time series better reflect underlying neural activity, increasing the sensitivity and interpretability of subsequent O-information analyses.

For ROI-level analysis, we used the 7-network parcellation of (Thomas Yeo et al. 2011), which defines seven canonical resting-state networks: visual (VN), somatomotor (SMN), dorsal attention (DAN), ventral attention (VAN), limbic (LIM), frontoparietal (FPN), and default mode network (DMN).

### Measuring functional HOI via the O-information

The time-series activity *X*(*t*) ≡ *X* from different N brain regions are represented as *X*_1_*, X*_2_*, …, X_N_*. The average information contained in those variables can be estimated via Shannon’s entropy *H*(*X*) = − ∑*_x_* prob(*x*)log(prob(*x*)), where *x* represents the possible states of variable *X* (Cover and Thomas 2005). The part of that information that is shared between pairs of variables *X*_1_ and *X*_2_ is measured by the mutual information *I*(*X*_1_*, X*_2_) = *H*(*X*_1_) + *H*(*X*_2_) − *H*(*X*_1_*, X*_2_), where *H*(*X*_1_*, X*_2_) is the joint entropy.

For three variables, the difference between pairwise and higher-order effects can be measured via the interaction information *II*(*X*_1_; *X*_2_; *X*_3_) = *I*(*X*_1_; *X*_3_) + *I*(*X*_2_; *X*_3_) − *I*(*X*_1_*, X*_2_; *X*_3_) (McGill 1954), where *I*(*X*_1_*, X*_2_; *X*_3_) is the mutual information between (*X*_1_*, X*_2_) and *X*_3_. Interestingly, *II*(*X*_1_; *X*_2_; *X*_3_) is symmetric in its three variables. For interactions involving more than three variables, the balance between low- and high-order effects can be captured via the O-information (Ω) (Rosas et al. 2019)

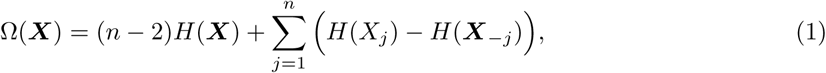

where ***X*** ≡ (*X*_1_*, X*_2_*, …, X_N_*) are the *n* random variables and ***X***_−_*_j_* ≡ (*X*_1_*, …, X_j_*_−1_*, X_j_*_+1_*, …, X_N_*) is equal to ***X*** with the component *X_j_* removed. Note that for *n* = 3 the O-information coincides with the interaction information. The sign of the O-information determines whether the interactions in the group ***X*** are dominated by redundancy (Ω(***X***) *>* 0) or synergy (Ω(***X***) *<* 0).

Although our analyses focus on triplets (n = 3), we adopt the O-information rather than McGill’s classical interaction information because it generalizes naturally to larger sets of variables. This provides a consistent formalism for quantifying redundancy and synergy across different group sizes, enabling straightforward extension to higher-order interactions beyond triplets. Building on this approach, for each subject, we calculate the O-information of triplets of variables 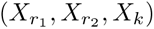, where 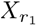 and 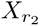 correspond to time series extracted from RSNs, rather than conventional ROIs, and *X_k_* is the time series of each voxel in the brain. Although the resulting O-information maps are reported at the voxel level, each voxel’s value reflects its contribution to interactions between RSNs, instead of indicating independent voxel-level network dynamics. As a result, for each participant we obtained 21 whole-brain O-information maps, one for each unique pairwise combination of the seven Yeo networks.

The O-information of each triplet was calculated by estimating the entropy using a rank-based normalization that provides a parsimonious estimate suitable for Gaussian processes (Ince et al. 2017). The code to compute the O-information is publicly available at (Marinazzo 2024).

### Conventional Neuroimaging Analysis

To assess group-level patterns and behavioral associations of high-order brain interactions, we defined two participant groups for statistical analysis. Group 1 consisted of 136 young adults (age *<* 30). Group 2 included 60 older adults (age *>* 60). The division of participants into “young” and “old” groups is artificial, as the age range in our sample is continuous (20-80 years). This grouping is used solely for visualization and descriptive purposes, rather than reflecting a true bimodal structure in the dataset. A third subgroup (Subgroup 1) was defined as a subset of Group 1 matched to Group 2 in terms of sex distribution, resulting in 60 participants. The same Subgroup 1 and Group 2 were used for behavioral associations. These analyses were conducted using Nilearn’s Second-Level Analysis framework (Abraham et al. 2014). Resulting statistical maps were thresholded using FDR correction at *p <* 0.05, with a cluster-size threshold of 50 voxels.

#### Group-level HOI

A voxel-level one-sample t-test on O-information maps within Group 1 was conducted. For each of the 21 ROI pair combinations, this analysis tested whether the mean O-information at each voxel significantly differed from zero across subjects, identifying brain regions that consistently exhibited redundancy- or synergy-dominated interactions with the corresponding ROI pair.

#### HOI associated with behavior

To examine the relationship between high-order brain interactions and cognitive performance, we used participants in Group 1. For each map, voxel-level GLMs were fitted with behavioral measures as independent variables. This approach allowed us to identify brain regions whose HOI significantly correlated with individual differences in behavior.

#### Group differences in HOI

To investigate age-related differences in high-order brain interactions, we performed voxel-level comparisons on the O-information maps. Specifically, a two-sample t-test was conducted using Group 2 (older adults) and Subgroup 1 (sex-matched young adults). We note that this division is artificial, as age in our sample spans a continuous range; the grouping primarily serves to highlight extremes of the age spectrum. This approach allowed us to identify brain regions showing apparent age-related differences in high-order statistical dependencies. From these regions, O-information values were extracted across voxels and subjects within each group. Distributions were then compared using the Mann–Whitney U test, and effect sizes (ES) were calculated. While this group-based analysis provides a descriptive view of age-related differences, complementary analyses treating age as a continuous variable would offer a more precise characterization of the relationship between age and high-order brain interactions.

#### Group differences in the association with behavior of HOI

To assess whether the relationship between behavioral performance and high-order brain interactions differed by age group (Group 2 vs Subgroup 1), we conducted a one-way ANCOVA covariate interaction (comparing regression coefficients between groups). For each O-information map, we computed the statistical contrast corresponding to the group × behavior interaction.

## Results

The methodological workflow is shown in Figure 1. Starting from a selected subgroup of participants from the preprocessed MPI-LEMON dataset (N = 196), the voxel-wise time series were preprocessed to mitigate physiological confounds (see Methods for details on RapidTide and rsHRF deconvolution). Subsequently, O-information was computed over triplets comprising two fixed region-level time series (defined here using canonical RSN masks) and a third signal corresponding to each voxel’s activity. This approach yielded whole-brain maps that quantified higher-order interactions, distinguishing redundancy (Ω *>* 0) and synergy (Ω *<* 0). These maps were then submitted to second-level voxelwise statistical analyses with corrections for multiple comparisons as specified previously, and finally used for conventional neuroimaging analysis.

**Figure 1:**
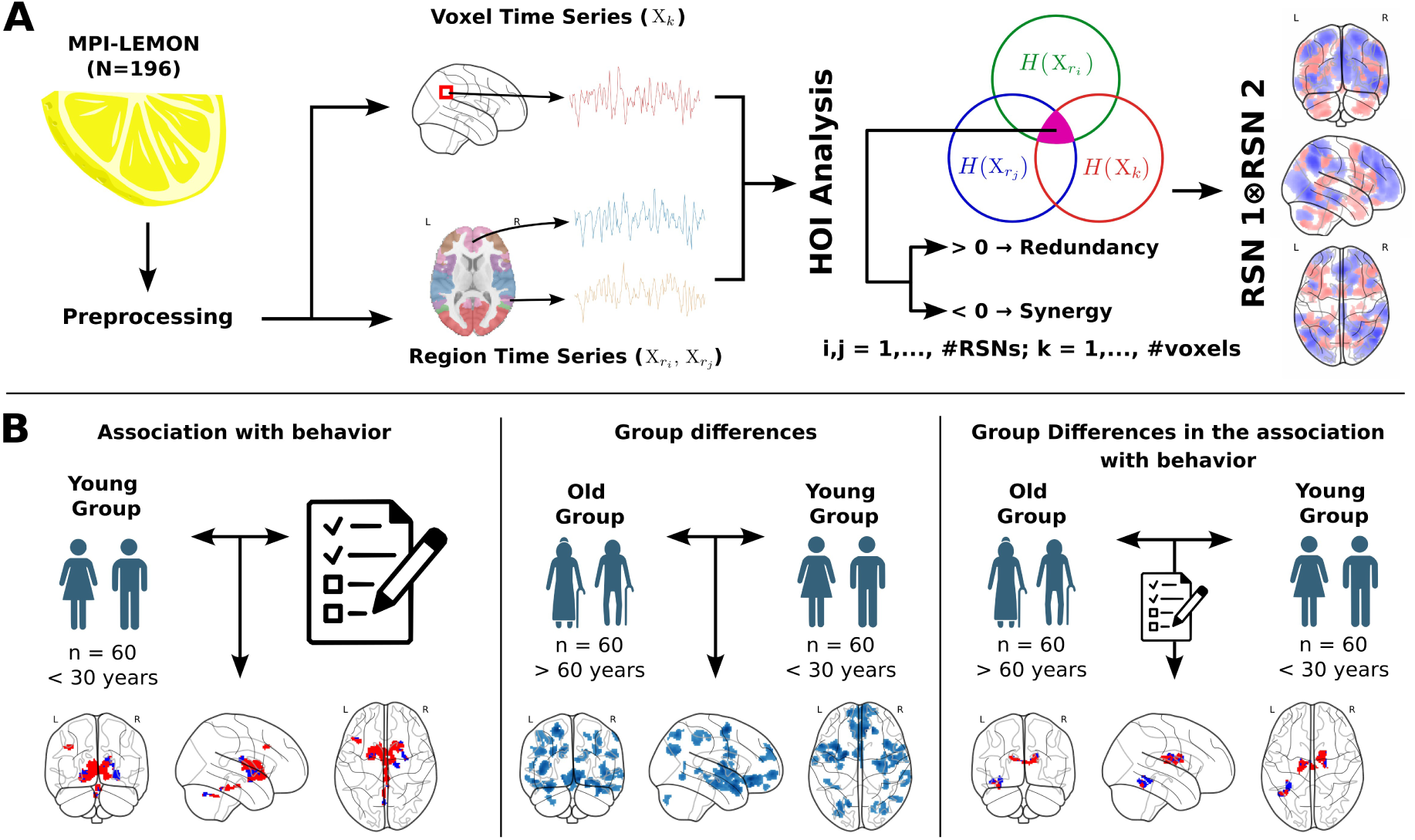
Methodological overview and workflow. **A:** A novel framework for voxel-level mapping of higher-order interactions in resting-state fMRI using O-information. Starting from preprocessed MPI-LEMON data (N = 196), the voxel-wise time series were preprocessed to mitigate physiological confounds (see Methods for details on RapidTide and rsHRF deconvolution). Region-level signals were computed as the mean time series within each anatomical region. O-information was then calculated over triplets consisting of two fixed region signals and a third variable of time series at voxel-level, thus the O-information reflects whole-brain maps that capture redundancy (Ω *>* 0) or synergy (Ω *<* 0) in these interactions. **B:** For showing feasibility, we conducted voxel-level statistical analyses to identify brain regions where O-information was associated with cognitive scores, differed by age, or showed age-dependent effects, after voxel-level multiple comparison correction as standard in conventional neuroimaging.

To characterize the spatial distribution of HOI, voxel-level one-sample t-tests were conducted on the O-information maps within Group 1. This analysis identified consistent patterns of redundancy and synergy across different combinations of RSNs (Figures 2a-b), and this depended on the specific network pairs included in each triplet. Overall, synergistic interactions were especially prominent in combinations involving the FPN, whereas redundancy tended to emerge in interactions involving other network pairs, such as the DMN or LIM. These findings indicate that distinct network pairings contribute differentially to higher-order functional organization in the resting brain.

**Figure 2a:**
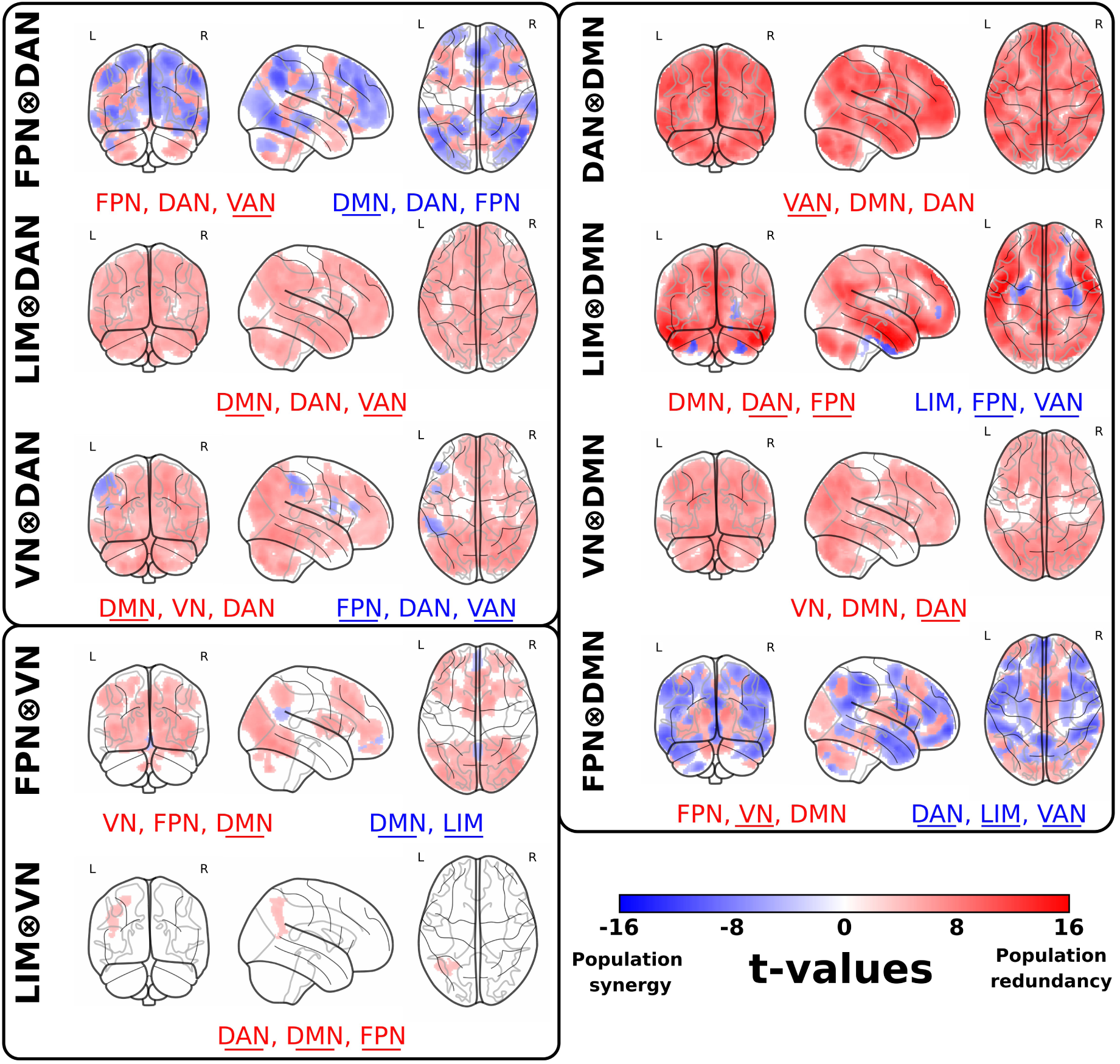
Brain maps of high-order functional interactions. Results of one-sample t-tests on O-information maps in helathy adults (Group 1), testing whether population-level O-information significantly differs from zero. O-information was computed for triplets consisting of two resting-state networks (RSNs, denoted by ⊗) and a third variable corresponding to any voxel in the brain. Positive t-values (red colors) indicate population-level redundancy, whereas negative t-values (blue colors) indicate population-level synergy. The figure is organized into three main panels, each fixing one reference RSN: the Dorsal Attention Network (DAN) (top left panel), the Visual Network (VN) (bottom left panel), and the Default Mode Network (DMN) (right panel). Beneath each set of brain maps, abbreviations of three RSNs are displayed. These correspond to the three networks whose spatial extents exhibit the greatest overlap with the significant t-values in that map. The color of each abbreviation reflects the predominant direction of the effect in the overlapping regions: red for positive t-values (redundancy), blue for negative t-values (synergy). Underlined abbreviations denote those RSNs different from the specific reference pair (i.e., the networks joined by the ⊗ symbol). L, left hemisphere; R, right hemisphere.

**Figure 2b:**
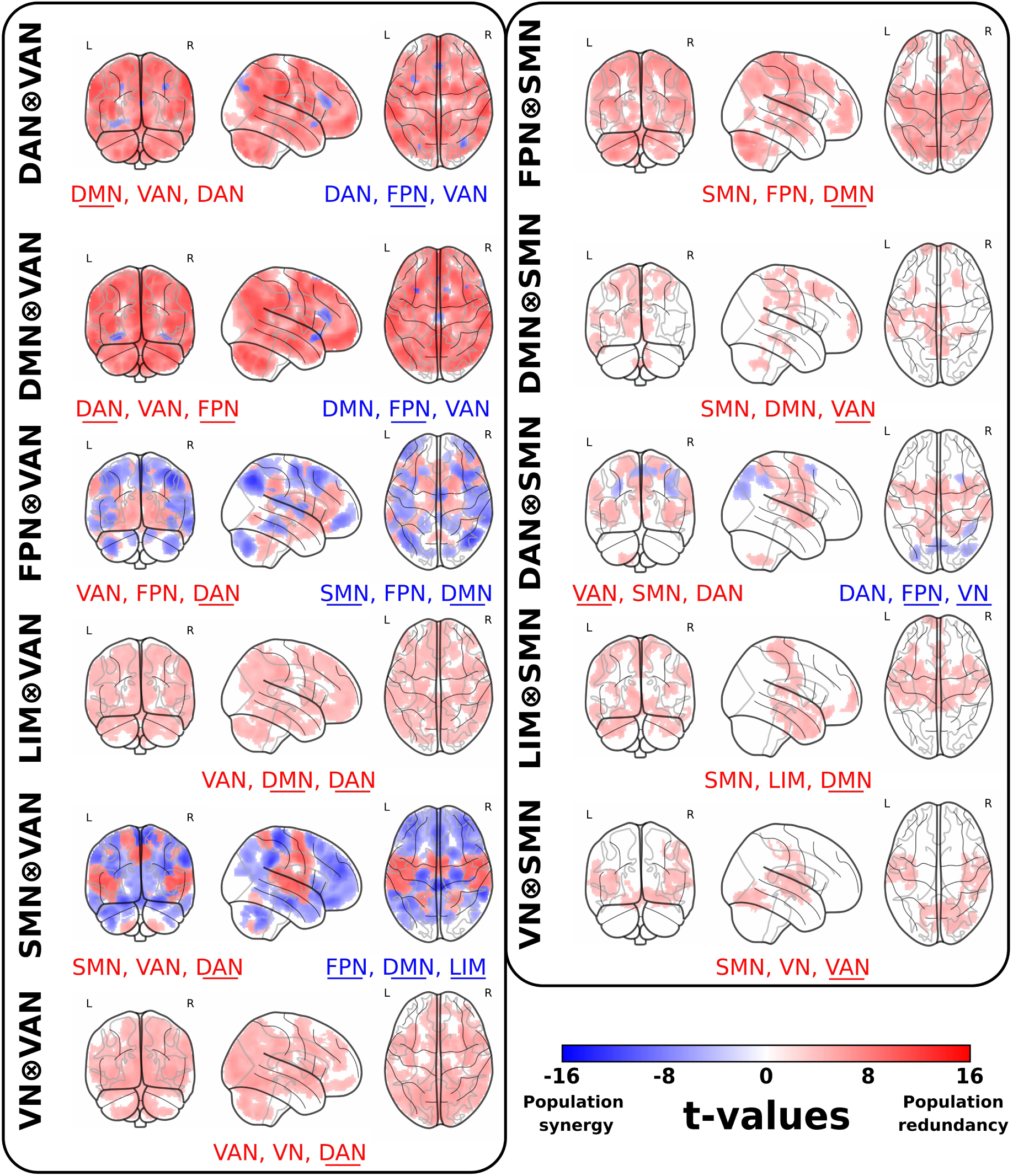
Brain maps of high-order functional interactions. Additional results as in Figure 2a., using the Ventral Attention Network (VAN) and Somatomotor Network (SMN) as reference RSNs.

To further characterize the anatomical distribution of redundancy- and synergy-dominated patterns observed at the population level (Figures 2a-b), we quantified the relative prevalence of synergistic and redundant voxels within each RSN. For each O-information map, voxels were classified according to the sign of Ω. For each individual RSN, we computed the percentage of synergistic and redundant voxels relative to the total number of voxels composing that RSN. Based on these quantities, we defined an Redundancy-Synergy balance (RSB) as:

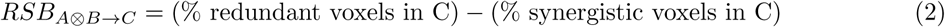

where *A* ⊗ *B* denotes the interacting RSN pair and *C* the RSN in which the voxel distribution is evaluated. Positive RSB values indicate a predominance of redundancy-dominated voxels within that RSN for the corresponding interaction pair, whereas negative values reflect a predominance of synergy-dominated voxels.

The resulting matrix (Figure 3) reveals a structured distribution of interaction regimes across large-scale networks. Several RSN combinations show predominantly positive RSB values within specific networks, indicating redundancy-dominated configurations, whereas others exhibit negative values consistent with a relative predominance of synergy. These patterns are not homogeneous across RSNs, suggesting that the spatial distribution of HOI depends on the particular combination of the interacting systems.

**Figure 3:**
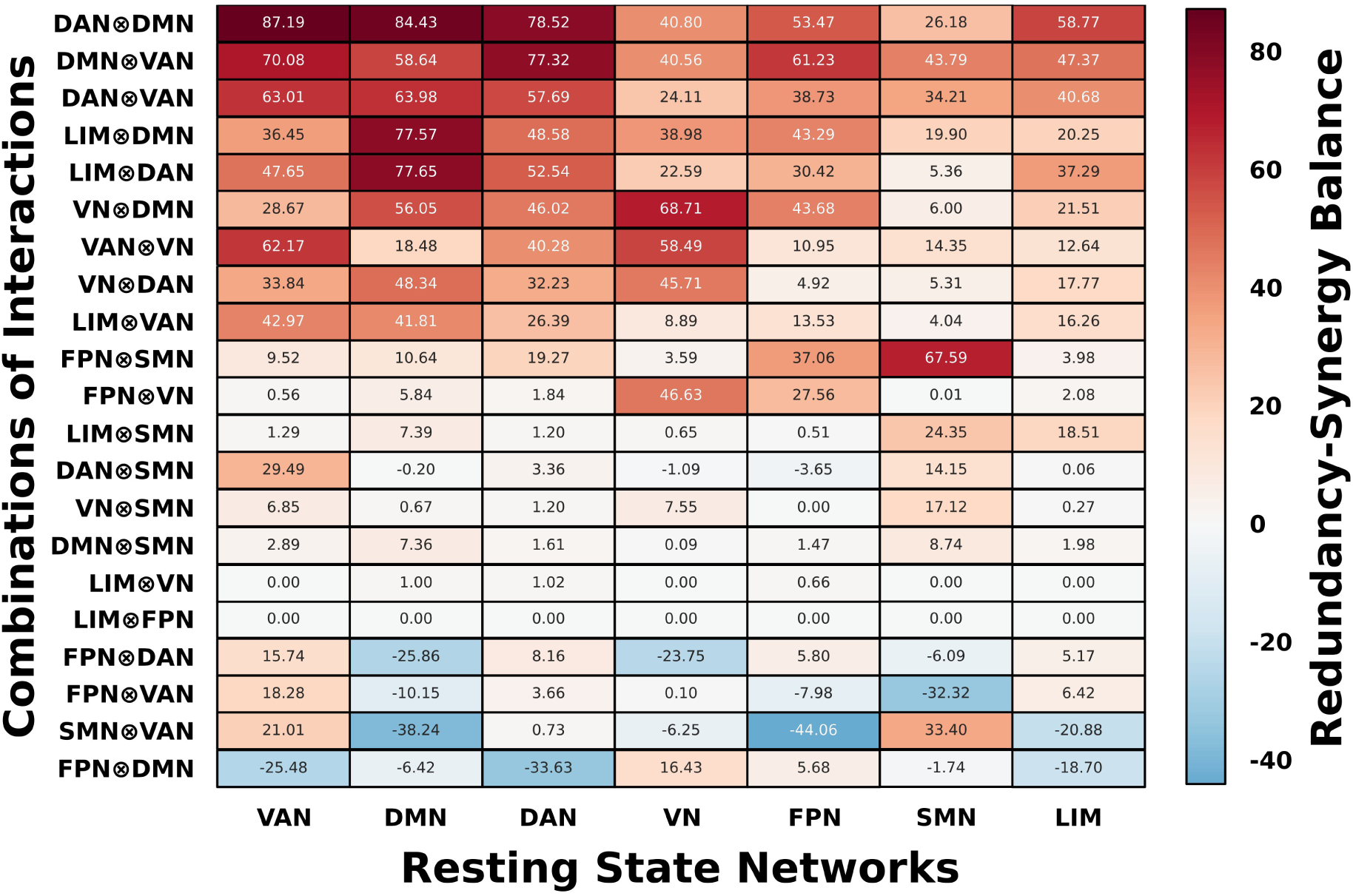
Network-level distribution of redundancy- and synergy-dominated voxels summarized by the Redundancy-Synergy balance (RSB). Matrix representation for each pair-wise combination of resting-state networks (rows) evaluated across individual resting-state networks (columns). For each RSN pair, and within each RSN, the percentage of redundant and synergistic voxels was computed relative to the total number of voxels composing that network. The RSB was defined as the difference between these quantities (RSB*_A_*_⊗_*_B_*_→_*_C_* = (% redundant voxels in resting state C) − (% synergistic voxels in resting state C)). Positive values indicate a predominance of redundancy-dominated voxels within the corresponding RSN, whereas negative values indicate a predominance of synergy-dominated voxels. This matrix provides a system-level summary of the spatial distribution of high-order interaction regimes across large-scale functional networks.

To examine the relationship between high-order brain interactions and cognitive performance, voxel-level GLMs were estimated using behavioral measures as the independent variable within Subgroup 1. Figure 4A shows the maps of significant voxels that survived FDR correction. Figure 4B displays, for those voxels, the median value of O-information across participants, colored in red when Ω *>* 0 and in blue when Ω *<* 0, corresponding respectively to redundancy and synergy. TMT B, assessing attention and processing speed, was correlated with both synergistic and redundant interactions in the FPN⊗DMN (t-values={min -6.32, max 4.93}, p-FDR*<*0.001). The RWT test, assessing verbal fluency and executive function, showed positive associations with redundant interactions in the VN⊗DMN (t-values={3.88, 5.16}, p-FDR*<*0.001). Performance in TAP VIS, assessing visual attention, was associated with synergistic interactions in the LIM⊗FPN (t-values={4.49, 5.29}, p-FDR*<*0.001), and with redundancy in the LIM⊗SMN interactions (t-values={4.09, 5.44}, p-FDR*<*0.001). In contrast, TAP WM, assessing working memory, was correlated with VN⊗SMN interactions (t-values={-6.69, -4.15}, p-FDR*<*0.001) in the opposite direction; higher synergy was associated with better performance. These results indicate that both redundancy and synergy in HOI contribute in a differente specific manner to task performance, particularly in the domains of attention, executive function, visual attention, and working memory.

**Figure 4:**
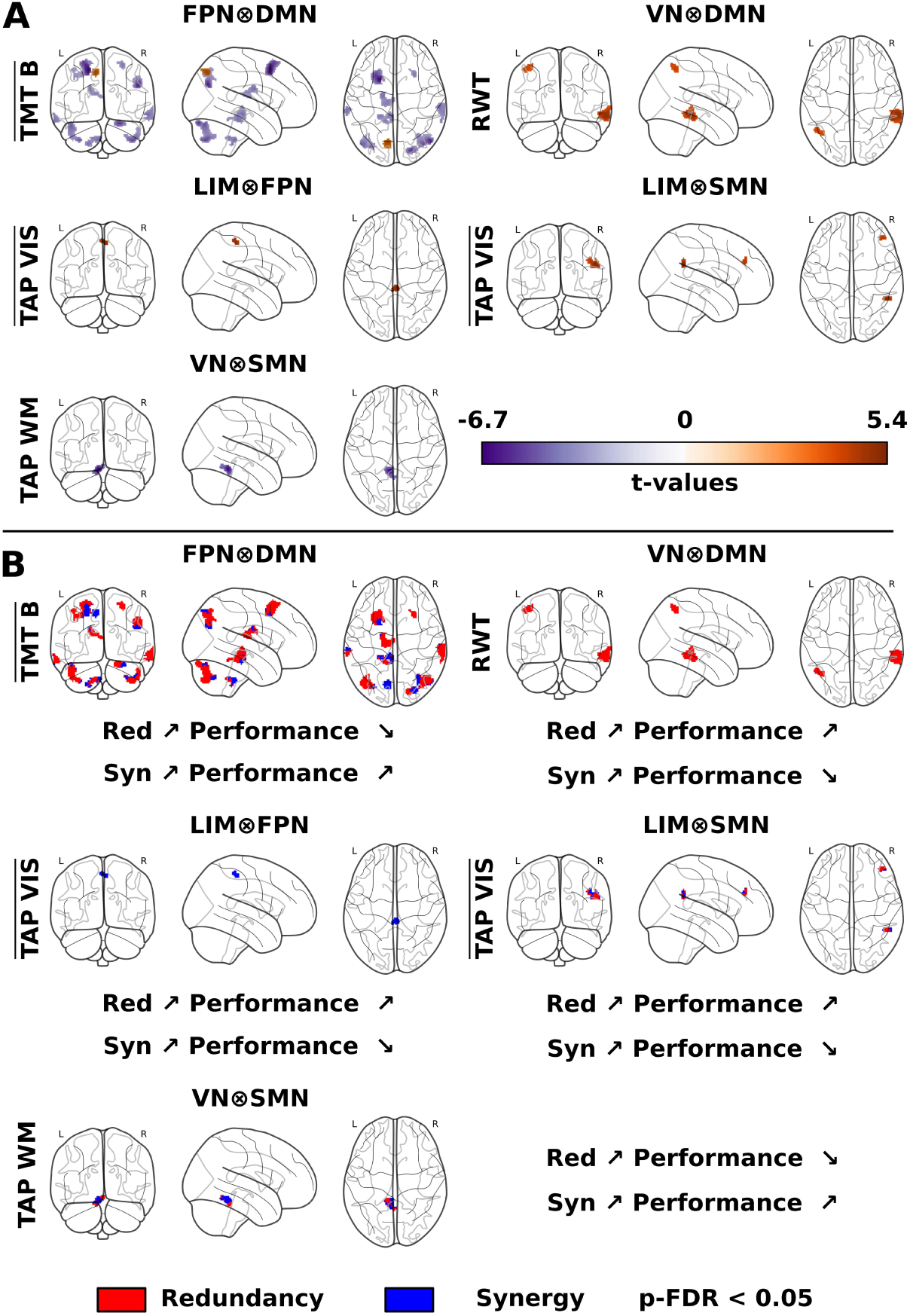
Relationship between high-order brain interactions and cognitive performance. (A) T-statistic maps from voxel-level GLM analyses showing brain regions where high-order interactions (HOI) between resting-state network (RSN) pairs were significantly associated with individual cognitive performance in young adults (Group 1), after FDR correction (p*<*0.05). Significant associations were found between TMT-B performance and FPN⊗DMN interactions, between RWT performance and VN⊗DMN interactions, between TAP VIS performance and both LIM⊗FPN and LIM⊗SMN interactions, and between TAP WM performance and VN⊗SMN interactions. (B) Median O-information value across participants are shown in red when Ω *>* 0 and in blue when Ω *<* 0, corresponding respectively to redundancy and synergy. Arrows below each map show the direction of association between redundancy or synergy and task performance (↗ increase, ↘ decrease). L, left hemisphere; R, right hemisphere.

To investigate how higher-order functional interactions differ across the lifespan, two-sample t-tests were conducted comparing Group 2 with Subgroup 1. Figure 5 highlights brain regions exhibiting significant age-related differences in O-information patterns. The most prominent changes were observed in DMN⊗DAN interactions (t-values={-5.24, -3.55}, p-FDR*<*0.001) (Figure 5A), where young adults displayed significantly greater redundancy than older adults (Mann-Whitney U: ES = -0.311). Likewise, DMN⊗LIM interactions (t-values={-5.02, -4.07}, p-FDR*<*0.001) revealed higher redundancy in young adults (Figure 5B) (Mann-Whitney U: ES = -0.369).

**Figure 5:**
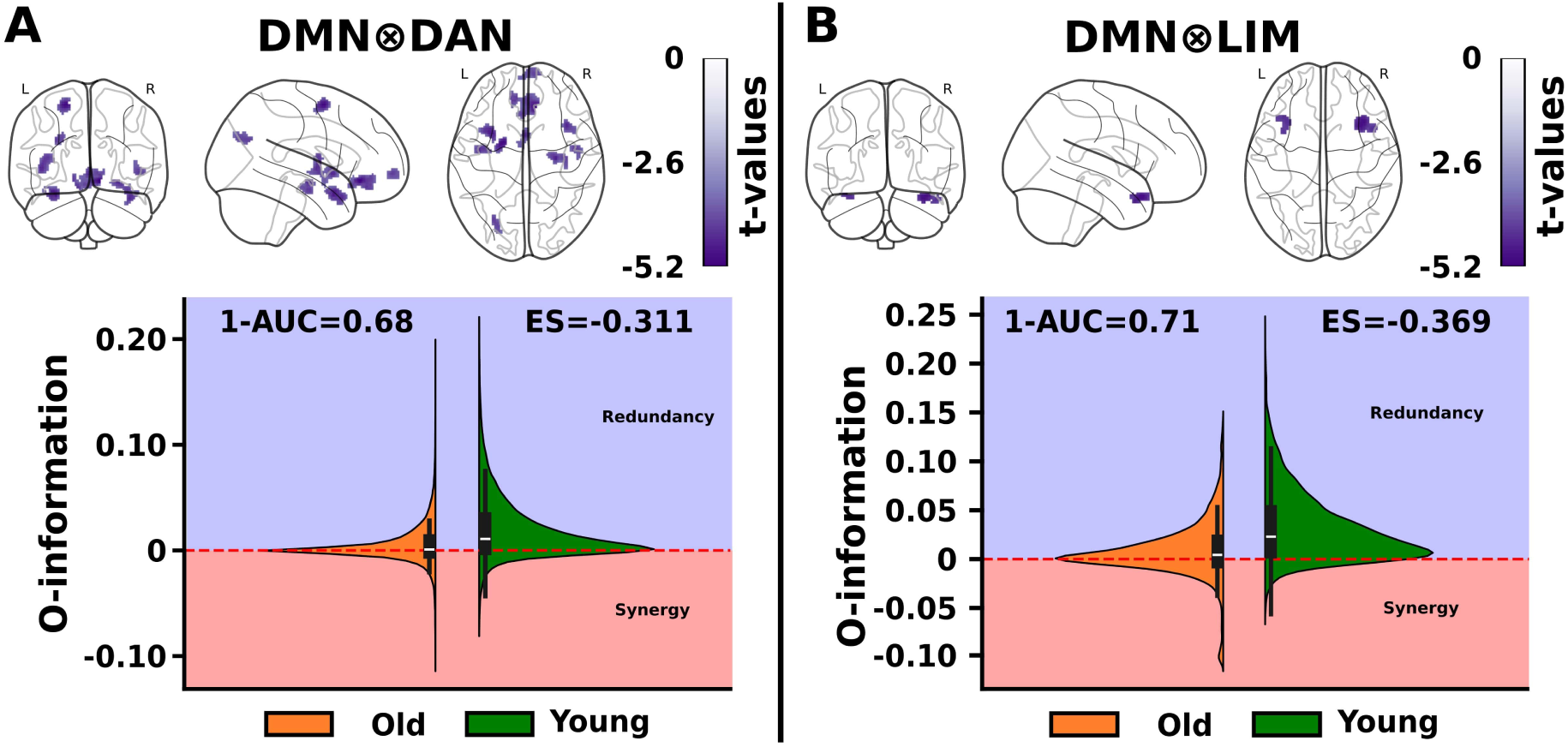
Group differences in high-order interactions. Results of two-sample t-tests comparing O-information maps between Group 2 and Subgroup 1 (old *>* young). Violin plots illustrate the distribution of O-information values across voxels and participants, with positive values (blue) indicating redundancy and negative values (red) indicating synergy. (A) DMN⊗DAN interactions showing significantly greater redundancy in young compared to older adults at the group level (Mann-Whitney U: ES = -0.311). (B) DMN⊗LIM interactions showing increased redundancy in young adults (Mann-Whitney U: ES = -0.369). ES: effect size. L, left hemisphere; R, right hemisphere.

To assess whether the relationship between higher-order interactions and behavioral performance varied across age groups, one-way ANCOVA analyses were conducted to examine group × behavior interactions between Group 2 and Subgroup 1. Figure 6A shows regions where the association between O-information and behavioral measures significantly differed between age groups. For both 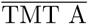, assessing attention and processing speed, and 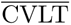, assessing episodic memory and verbal learning, negative associations with LIM⊗SMN (t-values={-6.19, -3.8}, pFDR*<*0.001) and LIM⊗VAN interactions (t-values={-5.52, -4.31}, pFDR*<*0.001), respectively, were observed between groups. To further interpret the group differences, we extracted the within-group t-maps and masked them using the significant voxels identified in the between-group contrast (Figure 6B). This approach allowed us to examine the direction and strength of behavior-related redundancy and synergy within each group, specifically in the regions driving the group-level differences. Additionally, to better characterize the underlying multivariate dependencies, we extracted the O-information values in these significant regions. For each group, we computed the median Ω across participants and visualized these values in Figure 6C.

**Figure 6:**
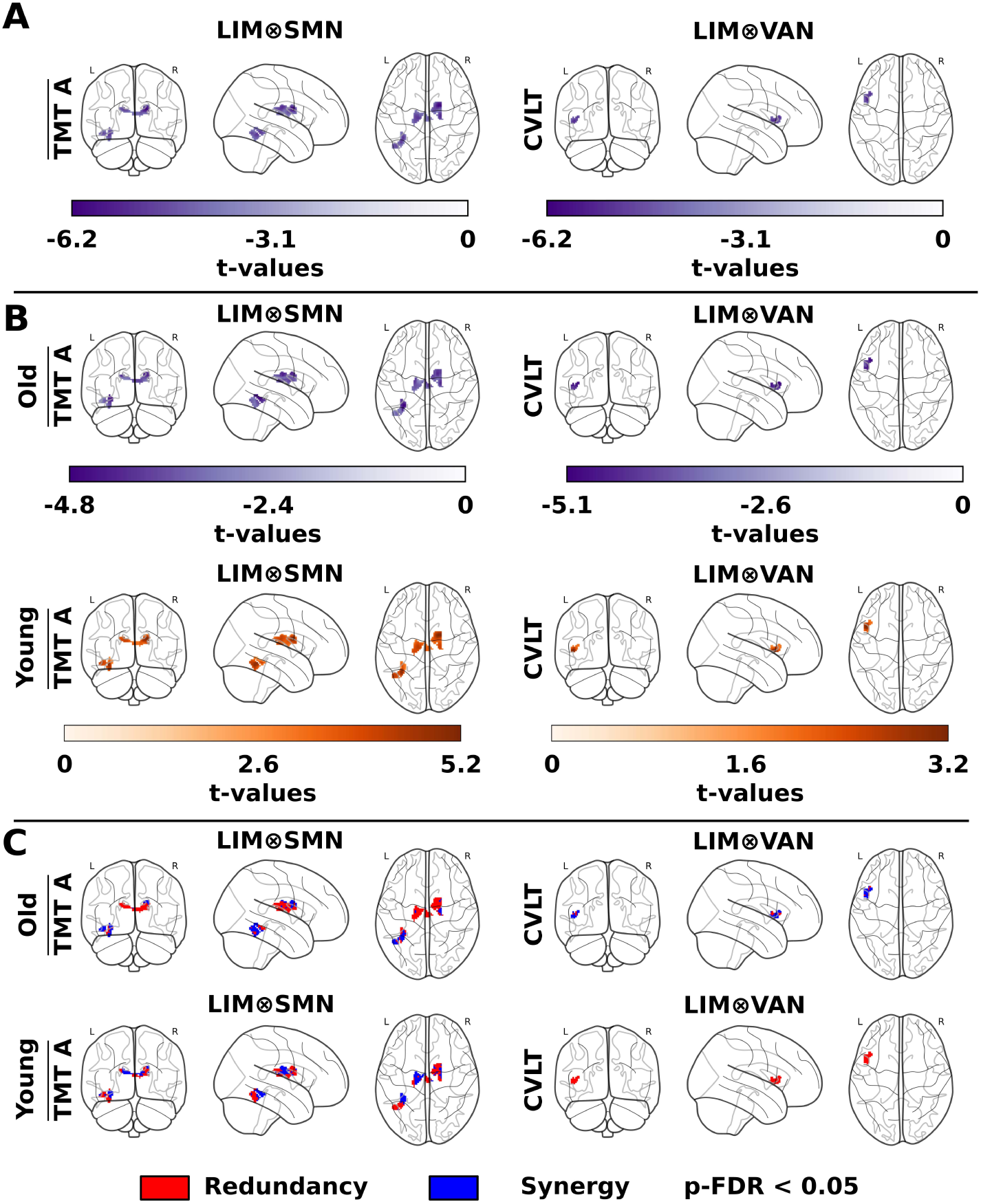
Group differences in the association with behavior of high-order interactions. Results of one-way ANCOVA covariate interactions (comparing regression between groups (Group 2 and Subgroup 1)). (A) Regions where the association between O-information and behavioral measures (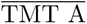 and 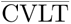) significantly differed between age groups, showing negative associations with LIM⊗SMN and LIM⊗VAN interactions, respectively. (B) Within-group t-maps, masked with significant voxels from the between-group contrast, illustrating the direction and strength of behavior-related redundancy and synergy in regions driving group-level differences. (C) Visualization of median O-information values across participants in significant regions, with voxels exhibiting Ω *>* 0 (redundancy-dominated, red) and Ω *<* 0 (synergy-dominated, blue), considering only those surviving FDR correction (p *<* 0.05). L left hemisphere; R, right hemisphere.

## Discussion

This study introduced a novel framework for integrating high-order functional interactions into conventional neuroimaging analysis using the O-information—a multivariate information-theoretic measure capable of distinguishing redundancy and synergy among n-plets of brain signals (here *n* = 3, but extension to higher *n* is straightforward). By applying this methodology to resting-state fMRI data, our analyses have shown voxel-level synergy and redudancy emerging at the population level, group differences between young and old adults and their association to cognitive performance, assessed by different neurophysicological tests.

A key step in our analysis is the construction of voxel-level HOI brain maps using triplets in which two variables correspond to two RSNs (we study any RSN pair interaction) and the third variable is the voxel time-series data. This strategy helps bridge the gap between traditional ROI-based network neuroscience and emerging high-dimensional information-theoretic frameworks. As a result, our framework supports full-brain statistical inference using conventional neuroimaging tools, enabling rigorous controls in the statistical analyses (e.g., FDR correction, cluster-size thresholding).

The population-level analysis of O-information maps revealed both redundant or synergistic voxel-level interactions. Synergistic interactions were especially prominent when the frontoparietal network was involved, while redundancy dominated in interactions recruiting the default mode and limbic networks. These results align with previous work showing a synergy-dominated frontoparietal network (Andrea I. Luppi et al. 2022) and a redundancy-dominated default mode network (Camino-Pontes et al. 2018). Notably, a synergistic role of the default mode network was also reported (Andrea I. Luppi et al. 2022). Differences arise from methodological distinctions between our approach and that of Luppi et al., who used Integrated Information Decomposition (Mediano et al. 2021) (ΦID) applied to time-delayed mutual information (TDMI). While both approaches aim to characterize the balance between synergy and redundancy, they differ in important ways. First, ΦID is based on TDMI, capturing dependencies between past and future states of neural signals, whereas the O-information quantifies instantaneous HOI across multiple variables. Second, ΦID requires solving an underdetermined system involving redundant, unique, and synergistic information components; in contrast, the O-information provides a single scalar value, making it more computationally tractable. Third, ΦID primarily focuses on pairwise systems evolving over time, while the O-information here was applied to triplets of variables. These results indicate that the specific mathematical formulation used to quantify synergy and redundancy plays an important role, and that the interpretability of these formulations must be considered with particular care.

To further summarize the spatial organization of HOI, we computed the balance between redundancy and synergy (RSB). We observed RSB patterns not uniformly distributed across the cortex, but instead following a network-dependent organization. Interactions involving the DMN, excluding the FPN (e.g., DAN⊗DMN, DMN⊗VAN), show consistently high redundancy values, whereas interactions involving the FPN or VAN with the SMN tend to shift toward synergy or reduced redundancy.

Our results indicate that specific HOI between large-scale RSNs are statiscaly significant related to individual differences in cognitive performance in young adults. Notably, synergy between the frontoparietal and default mode networks (FPN⊗DMN) showed a positive association with TMT B performance, suggesting that stronger synergistic integration between these networks may support more effective cognitive flexibility. In contrast, redundancy within the visual and somatomotor networks (VN⊗SMN) was positively related to working memory performance (TAP WM), highlighting the importance of stable and overlapping information processing in sensorimotor systems for memory function. Furthermore, visual attention (TAP VIS) was associated with distinct interaction patterns involving the limbic system, indicating that both redundant and synergistic relationships between limbic, frontoparietal, and sensorimotor networks may play a role in attentional control. These results support the notion that cognitive performance is not uniformly supported by one type of high-order interaction, but rather depends on a context-specific balance between redundancy and synergy across functional subsystems.

The study also examined age-related differences in HOI by comparing young and older adults, showing a pronounced reduction in redundancy-dominated DMN interactions in older adults, particularly in DMN⊗DAN and DMN⊗LIM pairs. This result suggests that aging may alter the capacity of the brain to sustain shared or overlapping informational dynamics across functionally distinct systems. This observations may be related to a decrease in network segregation found in aging brains (Pedersen et al. 2021; Zhang and Diaz 2023; Nolin et al. 2025). The higher redundancy observed in young adults may reflect a more consistent aligned functional coupling across DMN and other networks, supporting coherent and stable patterns of large-scale communication. In contrast, the reduced redundancy in older adults may indicate a loss of this shared informational structure, possibly reflecting reduced precision or coordination in functional dynamics. Note that studies examining interactions between individual brain regions and the rest of the brain, such as (Camino-Pontes et al. 2018), report increased redundancy in the DMN in older adults. This differs from the approach used here, where we focus on specific interactions between RSN pairs. In this framework, interactions involving the DMN—particularly DMN⊗DAN and DMN⊗LIM—tend to show reduced redundancy. These age-related differences may reflect broader changes in the efficiency and organization of high-order integration across the brain.

When looking at the relationship between HOI and behavior, negative associations between performance on TMT A and LIM⊗SMN interactions were found, as well as between CVLT and LIM⊗VAN, and these significant association was modulated by age. This suggests that young and older adults rely on different patterns of network interactions to support task performance. These findings are consistent with compensatory models of cognitive aging, where older brains may recruit alternative networks or adjust coordination strategies to maintain cognitive function despite changes in network organization (Heuninckx et al. 2008; Park and Reuter-Lorenz 2009; Goh and Park 2009; Sala-Llonch et al. 2015; Santos Monteiro et al. 2017; Rueda-Delgado et al. 2019). The within-group analyses indicate that older adults predominantly show negative associations between network interactions and cognitive performance, while young adults show positive associations in overlapping regions. The spatial distribution of these effects, particularly those involving limbic networks, may reflect age-related changes in emotional regulation and motivation that influence cognitive performance, aligned with results in (Bonifazi et al. 2018), where the fronto-striato-thalamic circuit was identified as showing a strong imprint of brain aging. These differential associations indicates that the relationship between redundancy-synergy patterns and behavior undergoes systematic modifications with aging.

A key limitation of the present framework lies in the necessity of dimensionality reduction to make the analysis of HOI tractable and interpretable. In our approach, we aimed to characterize the interaction between pairs of RSNs and the rest of the brain by forming triplets composed of two RSN time series and one voxelwise signal. However, each RSN is composed of thousands of voxels, needing a reduction to a single representative time series per network to construct these triplets. In this study, we chose dimensionality reduction by computing the mean time series within each RSN as a common and computationally simple strategy. Previous work has explored hyperharmonic representations to encode the full combinatorial structure of high-order signals (Medina-Mardones et al. 2021); however, these approaches are primarily designed for entire networks rather than for analyses that retain voxelwise contributions to RSN⊗RSN interactions. While alternative dimensionality reduction methods such as principal component analysis (PCA) could, in principle, offer more sophisticated representations, they have been shown to preferentially capture high-order redundancies while failing to preserve the full spectrum of multivariate dependencies, particularly higher-order synergies and topological structure (Varley, Mediano, et al. 2025). As such, the choice followed here represents a first step in this direction although we are aware that other methods could better capture the high dimensionality of the data.

A promising direction for future research is to examine how HOI patterns differ across various brain pathologies, potentially revealing novel biomarkers of dysfunction or compensation. Another important avenue involves the use of alternative methods for building brain maps using other metrics capturing HOI, such as information flow analysis (Andrea I. Luppi et al. 2022) or time-resolved measures of high-order dynamics (Varley, Pope, et al. 2023; Pope et al. 2024).

Overall, this study shows that high-order functional interactions, quantified using O-information, can be integrated into conventional neuroimaging analyses beyond their usual analytic framework. Our results support their value as biologically meaningful and reproducible informational markers, capturing both redundancy and synergy, and providing additional insight into behavior, aging, and brain disorders.

## References

Abraham, Alexandre, Fabian Pedregosa, Michael Eickenberg, Philippe Gervais, Andreas Mueller, Jean Kossaifi, Alexandre Gramfort, Bertrand Thirion, and Gaël Varoquaux (2014). “Machine Learning for Neuroimaging with Scikit-Learn”. In: Front. Neuroinform. 8.

Aschenrenner, Sonja, Oliver Tucha, and Klaus W. Lange (2000). Regensburger Wortflüssigkeits-Test: RWT. Verlag für Psychologie. Göttingen: Hogrefe.

Avants, Brian B., Nicholas J. Tustison, Gang Song, Philip A. Cook, Arno Klein, and James C. Gee (Feb. 2011). “A Reproducible Evaluation of ANTs Similarity Metric Performance in Brain Image Registration”. In: NeuroImage 54.3, pp. 2033–2044.

Babayan, Anahit et al. (2018). Max Planck Institut Leipzig Mind-Brain-Body Dataset - LEMON.

Babayan, Anahit, et al. (Feb. 2019). “A Mind-Brain-Body Dataset of MRI, EEG, Cognition, Emotion, and Peripheral Physiology in Young and Old Adults”. In: Sci Data 6.1, p. 180308.

Behzadi, Yashar, Khaled Restom, Joy Liau, and Thomas T. Liu (Aug. 2007). “A Component Based Noise Correction Method (CompCor) for BOLD and Perfusion Based fMRI”. In: NeuroImage 37.1, pp. 90–101.

Bonifazi, Paolo, Asier Erramuzpe, Ibai Diez, Iñigo Gabilondo, Matthieu P. Boisgontier, Lisa Pauwels, Sebastiano Stramaglia, Stephan P. Swinnen, and Jesus M. Cortes (Dec. 2018). “Structure–Function Multi-scale Connectomics Reveals a Major Role of the Fronto-striato-thalamic Circuit in Brain Aging”. In: Human Brain Mapping 39.12, pp. 4663–4677.

Bullmore, Ed and Olaf Sporns (Mar. 2009). “Complex Brain Networks: Graph Theoretical Analysis of Structural and Functional Systems”. In: Nat Rev Neurosci 10.3, pp. 186–198.

Camino-Pontes, Borja, Ibai Diez, Antonio Jimenez-Marin, Javier Rasero, Asier Erramuzpe, Paolo Bonifazi, Sebastiano Stramaglia, Stephan Swinnen, and Jesus Cortes (Sept. 2018). “Interaction Information along Lifespan of the Resting Brain Dynamics Reveals a Major Redundant Role of the Default Mode Network”. In: Entropy 20.10, p. 742.

Cover, Thomas M. and Joy A. Thomas (Sept. 2005). Elements of Information Theory. 1st ed. Wiley.

Cox, Robert W. (June 1996). “AFNI: Software for Analysis and Visualization of Functional Magnetic Resonance Neuroimages”. In: Computers and Biomedical Research 29.3, pp. 162–173.

Delis, Dean C., Joel H. Kramer, Edith Kaplan, and Beth A. Ober (Nov. 2016). California Verbal Learning Test–Second Edition.

Erramuzpe, Asier, Guillermo J Ortega, Jesus Pastor, Rafael G De Sola, Daniele Marinazzo, Sebastiano Stramaglia, and Jesus M Cortes (Dec. 2015). “Identification of Redundant and Synergetic Circuits in Triplets of Electrophysiological Data”. In: Journal of Neural Engineering 12.6, p. 066007.

Esteban, Oscar, Christopher J. Markiewicz, Ross W. Blair, Craig A. Moodie, A. Ilkay Isik, Asier Erramuzpe, James D. Kent, Mathias Goncalves, Elizabeth DuPre, Madeleine Snyder, Hiroyuki Oya, Satrajit S. Ghosh, Jessey Wright, Joke Durnez, Russell A. Poldrack, and Krzysztof J. Gorgolewski (Jan. 2019). “fMRIPrep: A Robust Preprocessing Pipeline for Functional MRI”. In: Nat Methods 16.1, pp. 111–116.

Fornito, Alex, Andrew Zalesky, and Edward T. Bullmore (2016). Fundamentals of Brain Network Analysis. San Diego: Elsevier Science & Technology.

Frederick, B. (2016-2024). rapidtide.

Friston, Karl J. (Jan. 1994). “Functional and Effective Connectivity in Neuroimaging: A Synthesis”. In: Human Brain Mapping 2.1, pp. 56–78.

Friston, Karl J. (2007). Statistical Parametric Mapping: The Analysis of Functional Brain Images. 1st ed. Amsterdam Boston: Elsevier / Academic Press.

Gatica, Marilyn, Cyril Atkinson-Clement, Carlos Coronel-Oliveros, Mohammad Alkhawashki, Pedro A. M. Mediano, Enzo Tagliazucchi, Fernando E. Rosas, Marcus Kaiser, and Giovanni Petri (Oct. 6, 2025). “Understanding the high-order network plasticity mechanisms of ultrasound neuromodulation”. In: PLOS Computational Biology 21.10. Ed. by Zhiyi Chen, e1013514.

Gatica, Marilyn, Rodrigo Cofré, Pedro A.M. Mediano, Fernando E. Rosas, Patricio Orio, Ibai Diez, Stephan P. Swinnen, and Jesus M. Cortes (Nov. 2021). “High-Order Interdependencies in the Aging Brain”. In: Brain Connectivity 11.9, pp. 734–744.

Gatica, Marilyn, Fernando E. Rosas, Pedro A. M. Mediano, Ibai Diez, Stephan P. Swinnen, Patricio Orio, Rodrigo Cofré, and Jesus M. Cortes (Sept. 2022). “High-Order Functional Redundancy in Ageing Explained via Alterations in the Connectome in a Whole-Brain Model”. In: PLoS Comput Biol 18.9, e1010431.

Goh, Joshua O. and Denise C. Park (Oct. 2009). “Neuroplasticity and Cognitive Aging: The Scaffolding Theory of Aging and Cognition”. In: Restorative Neurology and Neuroscience 27.5, pp. 391–403.

Greve, Douglas N. and Bruce Fischl (Oct. 2009). “Accurate and Robust Brain Image Alignment Using Boundary-Based Registration”. In: NeuroImage 48.1, pp. 63–72.

Herzog, Rubén, Fernando E. Rosas, Robert Whelan, Sol Fittipaldi, Hernando Santamaria-Garcia, Josephine Cruzat, Agustina Birba, Sebastian Moguilner, Enzo Tagliazucchi, Pavel Prado, and Agustin Ibanez (Dec. 2022). “Genuine High-Order Interactions in Brain Networks and Neurodegeneration”. In: Neurobiology of Disease 175, p. 105918.

Heuninckx, Sofie, Nicole Wenderoth, and Stephan P. Swinnen (Jan. 2008). “Systems Neuroplasticity in the Aging Brain: Recruiting Additional Neural Resources for Successful Motor Performance in Elderly Persons”. In: J. Neurosci. 28.1, pp. 91–99.

Ince, Robin A.A., Bruno L. Giordano, Christoph Kayser, Guillaume A. Rousselet, Joachim Gross, and Philippe G. Schyns (Mar. 2017). “A Statistical Framework for Neuroimaging Data Analysis Based on Mutual Information Estimated via a Gaussian Copula”. In: Human Brain Mapping 38.3, pp. 1541–1573.

Jenkinson, Mark, Peter Bannister, Michael Brady, and Stephen Smith (Oct. 2002). “Improved Optimization for the Robust and Accurate Linear Registration and Motion Correction of Brain Images”. In: NeuroImage 17.2, pp. 825–841.

Jenkinson, Mark, Christian F. Beckmann, Timothy E.J. Behrens, Mark W. Woolrich, and Stephen M. Smith (Aug. 2012). “FSL”. In: NeuroImage 62.2, pp. 782–790.

Korponay, Cole, Amy C. Janes, and Blaise B. Frederick (June 2024). “Brain-Wide Functional Connectivity Artifactually Inflates throughout Functional Magnetic Resonance Imaging Scans”. In: Nat Hum Behav 8.8, pp. 1568–1580.

Kumar G., Pradeep, Rajanikant Panda, Kanishka Sharma, A. Adarsh, Jitka Annen, Charlotte Martial, Marie-Elisabeth Faymonville, Steven Laureys, Corine Sombrun, Ramakrishnan Angarai Ganesan, Audrey Vanhaudenhuyse, and Olivia Gosseries (June 2024). “Changes in High-Order Interaction Measures of Synergy and Redundancy during Non-Ordinary States of Consciousness Induced by Meditation, Hypnosis, and Auto-Induced Cognitive Trance”. In: NeuroImage 293, p. 120623.

Leclercq, Michel and Peter Zimmermann, eds. (2002). Applied Neuropsychology of Attention: Theory, Diagnosis, and Rehabilitation. London New York: Psychology Press.

Li, Qiang, Vince D. Calhoun, Adithya Ram Ballem, Shujian Yu, Jesus Malo, and Armin Iraji (2023). Aberrant High-Order Dependencies in Schizophrenia Resting-State Functional MRI Networks.

Luppi, Andrea I, Pedro AM Mediano, Fernando E Rosas, Judith Allanson, John Pickard, Robin L Carhart-Harris, Guy B Williams, Michael M Craig, Paola Finoia, Adrian M Owen, Lorina Naci, David K Menon, Daniel Bor, and Emmanuel A Stamatakis (July 2024). “A Synergistic Workspace for Human Consciousness Revealed by Integrated Information Decomposition”. In: eLife 12, RP88173.

Luppi, Andrea I., Pedro A. M. Mediano, Fernando E. Rosas, Negin Holland, Tim D. Fryer, John T. O’Brien, James B. Rowe, David K. Menon, Daniel Bor, and Emmanuel A. Stamatakis (June 2022). “A Synergistic Core for Human Brain Evolution and Cognition”. In: Nature Neuroscience 25.6, pp. 771–782.

Marinazzo, Daniele (Mar. 2024). Github Repositiory for Retrieving High-Order Information Multiplets from Data Using the O-information.

McGill, William J. (June 1954). “Multivariate Information Transmission”. In: Psychometrika 19.2, pp. 97–116.

Mediano, Pedro A. M., Fernando E. Rosas, Andrea I Luppi, Robin L. Carhart-Harris, Daniel Bor, Anil K. Seth, and Adam B. Barrett (2021). Towards an Extended Taxonomy of Information Dynamics via Integrated Information Decomposition.

Medina-Mardones, Anibal M, Fernando E Rosas, Sebastián E Rodríguez, and Rodrigo Cofré (May 2021). “Hyperharmonic analysis for the study of high-order information-theoretic signals”. In: Journal of Physics: Complexity 2.3, p. 035009.

Mendes, Natacha, et al. (Feb. 2019). “A Functional Connectome Phenotyping Dataset Including Cognitive State and Personality Measures”. In: Sci Data 6.1, p. 180307.

Meunier, David, Renaud Lambiotte, and Edward T. Bullmore (2010). “Modular and Hierarchically Modular Organization of Brain Networks”. In: Front. Neurosci. 4.

Mirjebreili, Morteza, Josu Martinez De Aguirre Ibarreta, Daniele Marinazzo, and Laetitia Chauvière (May 2025). “High-Order Information Analysis of Epileptogenesis in the Pilocarpine Rat Model of Temporal Lobe Epilepsy”. In: eNeuro 12.5, ENEURO.0403–24.2025.

Nolin, Sara A, et al. (Mar. 2025). “Network Segregation Is Associated with Processing Speed in the Cognitively Healthy Oldest-Old”. In: eLife 14, e78076.

Park, Denise C. and Patricia Reuter-Lorenz (Jan. 2009). “The Adaptive Brain: Aging and Neurocognitive Scaffolding”. In: Annu. Rev. Psychol. 60.1, pp. 173–196.

Pedersen, Robin, Linda Geerligs, Micael Andersson, Tetiana Gorbach, Bárbara Avelar-Pereira, Anders Wåhlin, Anna Rieckmann, Lars Nyberg, and Alireza Salami (Nov. 2021). “When Functional Blurring Becomes Deleterious: Reduced System Segregation Is Associated with Less White Matter Integrity and Cognitive Decline in Aging”. In: NeuroImage 242, p. 118449.

Pope, Maria, Thomas F. Varley, and Olaf Sporns (2024). “Time-Varying Synergy/Redundancy Dominance in the Human Cerebral Cortex”. In.

Rangaprakash, D., Guo-Rong Wu, Daniele Marinazzo, Xiaoping Hu, and Gopikrishna Deshpande (Oct. 2018). “Hemodynamic Response Function (HRF) Variability Confounds Resting-state fMRI Functional Connectivity”. In: Magnetic Resonance in Med 80.4, pp. 1697–1713.

Reitan, Ralph (Oct. 1992). Neuropsychological Evaluation of Older Children.

Rokem, A., M. Trumpis, and F. Perez (2009). “Niftime: Time-Series Analysis for Neuroimaging Data.” In: Proceedings. of the 8th Python in Science Conference 2. Caltech, pp. 68–75.

Rosas, Fernando E., Pedro A. M. Mediano, Michael Gastpar, and Henrik J. Jensen (Sept. 2019). “Quantifying High-Order Interdependencies via Multivariate Extensions of the Mutual Information”. In: Physical Review E 100.3, p. 032305.

Rubinov, Mikail and Olaf Sporns (June 2011). “Weight-Conserving Characterization of Complex Functional Brain Networks”. In: NeuroImage 56.4, pp. 2068–2079.

Rueda-Delgado, Laura Milena, Kirstin Friederike Heise, Andreas Daffertshofer, Dante Mantini, and Stephan Patrick Swinnen (May 2019). “Age-Related Differences in Neural Spectral Power during Motor Learning”. In: Neurobiology of Aging 77, pp. 44–57.

Sala-Llonch, Roser, David Bartrés-Faz, and Carme Junqué (May 2015). “Reorganization of Brain Networks in Aging: A Review of Functional Connectivity Studies”. In: Front. Psychol. 6.

Santos Monteiro, Thiago, Iseult A.M. Beets, Matthieu P. Boisgontier, Jolien Gooijers, Lisa Pauwels, Sima Chalavi, Brad King, Geneviève Albouy, and Stephan P. Swinnen (Oct. 2017). “Relative Cortico-Subcortical Shift in Brain Activity but Preserved Training-Induced Neural Modulation in Older Adults during Bimanual Motor Learning”. In: Neurobiology of Aging 58, pp. 54–67.

Stramaglia, Sebastiano, Tomas Scagliarini, Bryan C. Daniels, and Daniele Marinazzo (Jan. 2021). “Quantifying Dynamical High-Order Interdependencies from the o-Information: An Application to Neural Spiking Dynamics”. In: Frontiers in Physiology 11, p. 595736.

Thomas Yeo, B. T., Fenna M. Krienen, Jorge Sepulcre, Mert R. Sabuncu, Danial Lashkari, Marisa Hollinshead, Joshua L. Roffman, Jordan W. Smoller, Lilla Zöllei, Jonathan R. Polimeni, Bruce Fischl, Hesheng Liu, and Randy L. Buckner (Sept. 2011). “The Organization of the Human Cerebral Cortex Estimated by Intrinsic Functional Connectivity”. In: Journal of Neurophysiology 106.3, pp. 1125–1165.

Varley, Thomas F., Pedro A. M. Mediano, Alice Patania, and Josh Bongard (2025). The Topology of Synergy: Linking Topological and Information-Theoretic Approaches to Higher-Order Interactions in Complex Systems.

Varley, Thomas F., Maria Pope, Maria Grazia, Joshua, and Olaf Sporns (July 2023). “Partial Entropy Decomposition Reveals Higher-Order Information Structures in Human Brain Activity”. In: Proceedings of the National Academy of Sciences 120.30, e2300888120.

Whitfield-Gabrieli, Susan and Alfonso Nieto-Castanon (June 2012). “Conn: A Functional Connectivity Toolbox for Correlated and Anticorrelated Brain Networks”. In: Brain Connectivity 2.3, pp. 125–141.

Wu, Guo-Rong, Nigel Colenbier, Sofie Van Den Bossche, Kenzo Clauw, Amogh Johri, Madhur Tandon, and Daniele Marinazzo (2021). “rsHRF: A Toolbox for Resting-State HRF Estimation and Deconvolution”. In: NeuroImage 244, p. 118591.

Zhang, Haoyun and Michele T. Diaz (June 2023). “Resting State Network Segregation Modulates Age-Related Differences in Language Production”. In: Neurobiology of Language 4.2, pp. 382–403.

